# Differences in microglia morphological profiles reflect divergent emotional temperaments: Insights from a selective breeding model

**DOI:** 10.1101/2021.05.28.445675

**Authors:** Pamela M. Maras, Peter Blandino, Elaine K. Hebda-Bauer, Stanley J. Watson, Huda Akil

**Affiliations:** The Molecular Neuroscience Institute, University of Michigan School of Medicine, Ann Arbor, MI 48109

**Author notes:** **Corresponding author:** Pamela M. Maras, Ph.D., University of Michigan School of Medicine, The Molecular Neuroscience Institute, 205 Zina Pitcher Place, Ann Arbor, MI 48109.

## Abstract

Microglia are known to play critical roles in healthy brain development and function, as well as the neuropathology underlying a range of brain diseases. Despite the growing evidence for a role for microglia in affective regulation and mood disorders, relatively little is known regarding how variation in microglia status relates to individual differences in emotionality. Using a selective breeding model based on locomotor response to novelty, we have generated rat lines with unique temperamental phenotypes that reflect broad emotional traits: low responder rats (bLRs) are novelty-averse and model a passive coping style, whereas high responder rats (bHRs) are highly exploratory and model an active coping style. To identify a possible functional role of microglia in these phenotypes, we administered minocycline, an antibiotic with potent microglia inhibiting properties. We found changes in emotional and social behaviors in bLRs, with no discernable effects in bHRs. Using detailed anatomical analyses, we went on to explore the nature of baseline differences in hippocampal microglia populations associated with the divergent temperaments. Interestingly, we found that although bHRs and bLRs had comparable total numbers of hippocampal microglia, selective breeding was associated with a shift in the morphological features of these cells. Specifically, microglia from bLRs were characterized by a hyper-ramified morphology, with longer processes and more complicated branching patterns than microglia from bHRs. This morphology is thought to reflect an early stage of microglia activation and suggests that microglia from bLRs may be in a reactive state even when animals are not overtly stressed or challenged. Taken together, our results provide novel evidence linking variation in inborn temperament with differences in the baseline morphological status of microglia cells and highlight the importance of considering the role of microglia not only in acute responses to stress but also in shaping enduring characteristics of emotionality.

## Introduction

The expression of emotional behaviors reflects complex interactions between genetic and environmental factors. Within the range of inborn factors, it is becoming increasingly recognized that broad personality traits represent a key predictor for emotional resilience or vulnerability (Geyer et al., 2010; Hink et al., 2013; Turner et al., 2017). Often referred to as temperament, considerable variation exists in the way individuals interact with their environment and respond to challenges (Clauss and Blackford, 2012; Clauss et al., 2015). Emotional temperaments represent traits that are highly stable throughout the lifespan and across different conditions, and research indicates that underlying temperament may be a risk factor for developing certain psychopathologies (Hink et al., 2013; Clauss et al., 2015). Indeed, individuals with a timid or novelty-averse temperament appear particularly vulnerable and are at higher risk for developing a range of psychiatric disorders, including clinical anxiety and depression (Caspi et al., 1996; Gladstone et al., 2005; Gladstone and Parker, 2006; Clauss and Blackford, 2012). Despite the recognition of temperament as a critical individual variable, the neurobiological mechanisms that underlie emotionality traits remain unclear.

Selective breeding models provide a useful tool to study the neurobiology of temperament and its relationship to emotional output. By selectively breeding rats based on locomotor response to novelty, our laboratory has generated lines with opposing temperaments: high responders (bHRs) and low responders (bLRs) (Stead et al., 2006). The bHRs are highly exploratory with an active coping style that models an “externalizing temperament”. By contrast, the bLRs are novelty-averse and represent a passive or inhibited coping style that models and “internalizing temperament” (Stead et al., 2006; Turner et al., 2017). These locomotor phenotypes are highly heritable (Zhou et al., 2019) and predict a wide range of emotional behaviors (Clinton et al., 2007, 2010, 2011b; Turner et al., 2008, 2011; Stedenfeld et al., 2011), as well as responsivity to pharmacological and environmental interventions (Jama et al., 2008; Perez et al., 2009; Calvo et al., 2011; Clinton et al., 2011a; Turner et al., 2011; Aydin et al., 2015; Rana et al., 2015). In multiple laboratory assays, bLRs display increased anxiety- and depressive-like behaviors and reduced social exploration and proactive coping, consistent within rodent models of internalizing disorders (Turner et al., 2017). Much research has focused on the neurobiology underlying these temperaments, and a recent meta-analysis of multiple transcriptional studies has identified broadly divergent patterns of gene expression within hippocampus, highlighting a range of cellular processes (Birt et al., 2020). Interestingly, microglia-related pathways emerged as one of the main functional candidates from this analysis, as the bLR phenotype is associated with elevated expression of several key genes related to microglial activation and signaling (Birt et al., 2020).

These findings are intriguing, as they complement a growing body of research implicating microglia in the development of mood disorders. Beyond serving as the brain’s innate immune cells, microglia play a critical role in the healthy development and dynamic functioning of the brain, regulating processes from cell survival to synaptic stability (Tremblay et al., 2011; Salter and Beggs, 2014; Schafer and Stevens, 2015). Microglia-related impairments appear to underlie some of the pathological changes associated with psychiatric diseases, including depression (Yirmiya et al., 2015; Mondelli et al., 2017). As a result, a neuroimmune hypothesis has emerged that suggests microglia dysfunction constitutes a potential risk factor or biomarker for these disorders (Yirmiya et al., 2015; Bhattacharya and Drevets, 2017; Mondelli et al., 2017). A neuroimmune hypothesis is supported by data from multiple rodent models that induce repeated or sustained stress, produce changes in depressive-like behaviors and show an association with alterations in microglia number and/or activity throughout the brain (reviewed in Delpech et al., 2015; Yirmiya et al., 2015). Moreover, the potent microglia inhibitor minocycline has been shown to mitigate many of the stress-induced behavioral deficits in these models, as well as some of the underlying microglia alterations (Henry et al., 2008; Molina-Hernández et al., 2008; Pae et al., 2008; O’Connor et al., 2009; Arakawa et al., 2012; Hinwood et al., 2012, 2013; Zhu et al., 2014; Kreisel et al., 2014; Zheng et al., 2015; Majidi et al., 2016; Tong et al., 2017; Wang et al., 2018; Zhang et al., 2018, 2019b, 2019a).

Taken together, these studies suggest that microglia play a role in the pathophysiology of affective disorders, yet critical questions remain regarding how individual differences in microglia relate to underlying temperament. One possibility was that microglia would show differential responses between individuals primarily under stressful conditions. However, the fact that our gene expression profiling was carried in unstressed animals led us to the hypothesis the bLR phenotype may be due, at least in part, to an over-activation of microglia even under basal conditions, implying a genetic role in the reactivity of microglia. To address this question, we administered minocycline and characterized its impact on the emotional and social behaviors of the two selectively bred lines. The differential effect of this drug on the two lines led us to a careful analysis of the baseline status of microglia as a function of selective breeding for differences in emotionality.

## Materials and Methods

### Animals

All subjects were male Sprague Dawley rats acquired from our in-house breeding colony at the Molecular and Behavioral Neuroscience Institute (spanning generations 52 – 65). The initial generation and maintenance of our breeding lines is described in detail elsewhere (Stead et al., 2006) and in *Supplemental Materials*. Briefly, adult rats were originally screened for their locomotor response to a novel environment, and the top and bottom 20% of responders were used to breed bHR and bLR lines, respectively. These lines have been maintained over several generations, resulting in highly stable and predictable divergent lines. All experimental procedures were approved by the University Committee on the Use and Care of Animals at the University of Michigan and were conducted in accordance with the National Institute of Health (NIH) Guide for the Care and Use of Laboratory Animals (2011).

### Quantitative reverse transcription-PCR (q-PCR)

To determine baseline expression differences, whole hippocampus was collected from adult bHRs and bLRs (*n* = 10 per line) under non-stress conditions. RNA was extracted using the RNeasy Mini Kit (Qiagen #74104), and cDNA was synthesized using iScript cDNA Synthesis Kit (Biorad #1708891). Amplification reactions were performed using a BioRadiCycler with SYBR Green detection. Samples were run in duplicate, and group differences in mean quantification cycle (Cq) values were calculated using the ΔΔCq method (Livak and Schmittgen, 2001), with glyceraldehyde-3-phosphate dehydrogenase (*Gapdh,* NM_017008) as the reference gene. See *Supplemental Materials* for full qPCR experiment details, validation procedures, and primer sequences (Table S1).

### Minocycline administration

To test the functional role of microglia on temperament, we used minocycline (Sigma Aldrich, St. Louis, MO) administration to inhibit microglia activity. To reduce handling, minocycline was dissolved in the drinking water (1mg/ml, pH adjusted to ~7.4) to achieve an estimated daily dose of 60-100 mg/kg. Minocycline (or tap water for controls) was administered in the drinking bottles for 14 days prior to the initiation of behavioral testing (15^th^ day, see below) and was continued throughout the duration of the experiment. The treatment regimen was chosen based on previous studies showing behavioral effects using similar doses and timing (Hinwood et al., 2012, 2013). Minocycline/water consumption was measured daily, and body weight was tracked over the course of the experiment. To confirm minocycline dosing, circulating levels of minocycline were measured from plasma collected in a sub-set of subjects at the end of the experiment using a commercially available tetracycline enzyme immunoassay (Perkin Elmer, Akron, OH, Part #FOOD-1016-04C; Hinwood et al., 2012) following kit instructions.

### Behavioral testing

For the first behavioral experiment, the effects of minocycline administration on the bLR phenotype were assessed using an array of behavioral tests. Following 2 weeks of minocycline or control water treatment (*n* = 10 per treatment group), bLRs were tested in the following sequence: novelty suppressed feeding, open field, social interaction, sucrose preference, and forced swim. To determine whether any behavioral effects were specific to the underlying temperament, a separate study was performed using the same minocycline regimen in both bHRs and bLRs (*n* = 8 per treatment group, per line), analyzing one anxiety-like measure (elevated plus maze) and one depressive-like measure (forced swim). All behavioral tests were conducted between 9:00 am and 1:00 pm and sequential tests were separated by 24 – 72 hours. All behavioral coding was done by a researcher who was blind to the identity of the subject. See *Supplemental Materials* for full details of each behavioral test.

### Iba1 immunohistochemistry

Adult rats (*n* = 8 per line) were transcardially perfused for brain tissue fixation with minimal disturbance (*Supplemental Materials*). Coronal brain sections (40-μm thickness) were collected using a cryostat (−20°C) in a 1:6 series, and sections were stored in cryoprotectant-antifreeze solution until processing. For processing, a single, randomly-selected series of sections from each subject were removed from cryoprotectant and processed using standard, free-floating immunohistochemical procedures (Hoffman et al., 2008; *Supplemental Materials*). Briefly, microglia cells were labeled using a primary antibody raised against the ionized calcium binding adaptor molecule 1 (Iba1, FUJIFILM Wako Chemicals, Richmond, VA, concentration 1:70,000) and visualized with nickel-enhanced diaminobenzidine (DAB) reaction. Iba1 is a microglia-specific marker that stains the cell body as well as the full extension of cell processes (Imai and Kohsaka, 2002), allowing analysis of key morphological features known to reflect the functional status of microglia cells (Kettenmann et al., 2011).

Iba1 staining was analyzed in multiple ways, described below. For all microscope analyses, live images were analyzed with a Leica DMR microscope attached to a digital camera and a three-axis motorized stage. All slides were coded, such that the researcher performing the analysis was blind to subject identity/group.

### Iba1 staining density

As an initial measurement of microglia, we calculated the optical density of Iba1 immunostaining in bHRs (*n* = 8) and bLRs (*n* = 7). Slides were scanned (PathScan Enabler; Meyer Instruments) and converted into 16-bit grayscale files. ImageJ Software (NIH) was used to quantify optical density of Iba-1 staining. The rostral hippocampal region was outlined in four consecutive sections per subject (Swanson, 2004; Atlas levels 28 – 34; Figure 1A). A threshold function was applied (Moments function), and the number of pixels surpassing threshold was calculated as a percentage of the region outlined. This method has been used to capture microglia soma and processes and reflect density of Iba1 signal (Perkins et al., 2018). Density values were averaged across all sections to generate a single value per subject.

**Figure 1.**
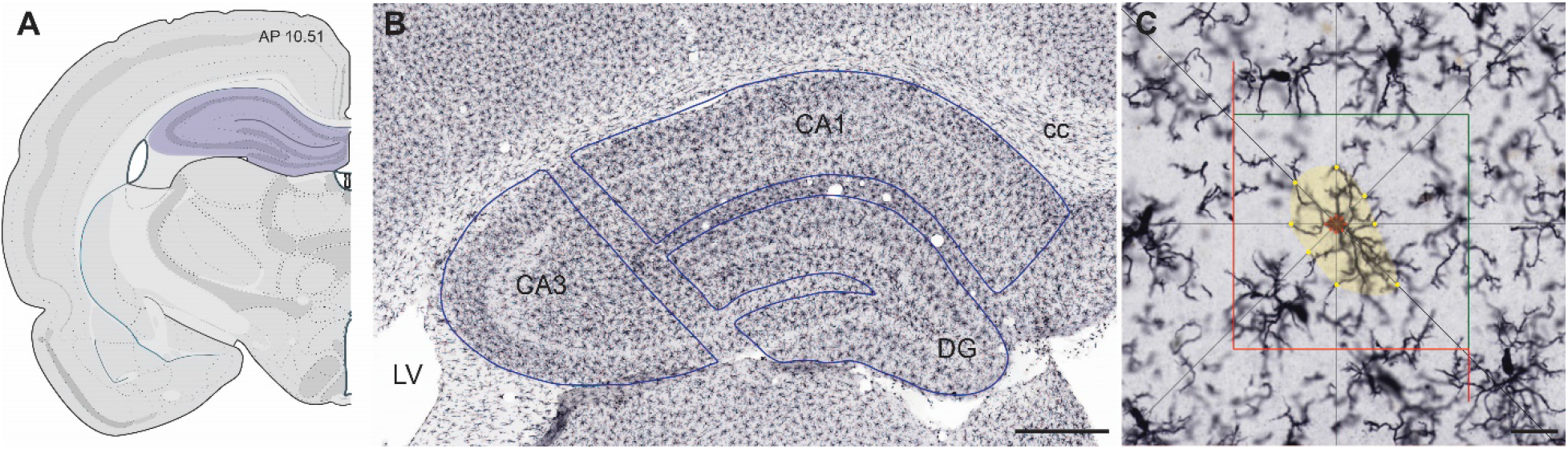
Stereological estimates of microglia number and area in hippocampus of bHRs and bLRs. (**A**) Coronal sections of dorsal hippocampus were matched for rostro-caudal level using a Rat Brain Atlas (Swanson, 2004; Atlas levels 28 – 34). (**B**) Photomicrograph showing immunohistochemical staining of microglia marker Iba-1. The different sub-regions of hippocampus were sampled separately during stereological sampling and were drawn using anatomical landmarks (blue lines). Scale bar = 500 μm. (**C**) Recreated depiction of workflow using Optical fractionator and Nucleator probes. Cells were counted if soma was within the sampling grid (red and green lines). When a cell was selected, the Nucleator probe applied 8 rays from the center of the cell, and points of intersection were used to estimate soma area (red markers) and cell territory (yellow markers, distal extensions of cell processes). Scale bar = 20 μm. Abbreviations: CA1 and CA3 = Cornu Ammonis 1 and 3; cc = corpus callosum; DG = dentate gyrus; LV = lateral ventricles.

### Microglia number

To compare the total numbers of microglia cells, non-biased stereological quantification was performed using the Optical Fractionator probe within Stereo Investigator software (MicroBrightField Bioscience, Williston, VT). For each subject (*n* = 7 per line), four sections within hippocampus were analyzed unilaterally. The Cornu Ammonis 1 and 3 (CA1 and CA3) and dentate gyrus (DG) sub-regions of hippocampus were outlined according to anatomical landmarks using a 2.5X objective and were analyzed separately (Figure 1B). Cells were counted using a 40X dry-objective lens. Stereological sampling parameters (Table S2) were determined through pilot studies to ensure sufficient sampling and Gundersen-Jensen error coefficients below 0.10 for all subjects. Iba1-positive cells were identified as having dark, blue–black somatic staining, and were excluded from counting if their cell body contacted the left or bottom edges of the counting frame (Figure 1C; West et al., 1991). A categorical assessment of microglia state (Kettenmann et al., 2011) was assigned to each cell while counting: *Ramified*: small cell body, with long/thin processes; *Reactive*: larger cell body, with few, stout processes; and *Amoeboid*: enlarged, round cell body, with minimal/no processes.

### Microglia area

The size of microglia cells, both the soma and its processes, is dynamic, and shifts in cell area can provide insight into the functional state of the cell, as well as quantitative information regarding variability of cells within a particular state (Kettenmann et al., 2011). To capture potential size differences in microglia cells across the hippocampal population, we generated area estimates for every cell counted during the stereological sampling procedure with the addition of the Nucleator probe (isotropic, 8-rays) (Gundersen, 1988; Perkins et al., 2018). As each cell was selected for counting, a ray of 8 lines was applied from the center of the cell (Figure 1C). Points of intersection along the rays were used to generate 2 area measurements: 1) intersections with the edge of the cell body were used to estimate *soma area* and 2) intersections with the most distal extent of the cell processes were used to generate overall *territory area*. In this way, we generated quantitative estimates of microglia cell area in a population-wide (hundreds of cells per subject) and non-biased manner.

### Microglia morphology

To explore the morphological characteristics of microglia cells in more detail, we generated full reconstructions of a subset of ramified microglia from bHRs and bLRs using Neurolucida software (MicroBrightField Bioscience). Cells were selected from a systematically placed circular region of interest (ROI) within CA1, CA3, and DG on a single section for each subject (*n* = 8 per line). Five cells were selected within each ROI, for a total of 40 cells per ROI, per line. Individual ramified cells were selected based on staining criteria that included a visible full cell body that was reasonably distinguishable from nearby cells, with no major disconnections of processes. Tracings were performed using a 63X-oil objective (1.32 numerical aperture) by first tracing the edge of the cell body at a single plane of focus. Each process was then traced, with both the width of the process and plane of focus adjusted incrementally. Tracing and editing continued until all visible processes were reconstructed. Reconstructions were analyzed using Neurolucida Explorer Software (MicroBrightField Bioscience), which generated multiple measurements for each cell drawn, including: soma area; total number, length, and surface area of processes, and total number of branches (nodes). Convex hull analysis, which calculates the area of a polygon joining the distal points of each of the cell processes, was also used to reflect the territory of each reconstructed cell (Hinwood et al., 2013). During the data processing step, a total of 4 cells (2 from bHRs; 2 from bLRs) were identified as extreme statistical outliers in multiple measures, indicating likely errors during the reconstruction process. These cells were removed from the analyses, leaving a final *n* = 38 cells per line.

### Statistical analysis

All data were analyzed with SPSS version 26 (IBM, Armonk, NY). For qPCR data, between group comparisons in -ΔΔCq values were made using Welch’s independent *t*-tests, due to unequal variance. For the minocycline study in bLRs only, effects of minocycline administration were determined using independent *t*-tests comparing minocycline- and water-treated groups. For the minocycline study comparing bHRs and bLRs, 2-way ANOVAs were performed with breeding line *X* minocycline treatment as between-subjects factors. Any significant interactions were explained using simple main effects. For Iba1 analyses, data were first analyzed with mixed-factors ANOVAs to determine if the effects of breeding line (bHR, bLR) varied across sub-region analyzed (CA1, CA3, DG). For data sets in which there was not a significant interaction, data were collapsed across sub-regions prior to final analysis and presentation. For data sets in which there were variations across sub-regions, results were analyzed and presented separately for each sub-region. Line differences in Iba1 measures were determined using independent samples *t*-tests, with bonferroni alpha corrections where appropriate. Full statistical values are presented in *Supplemental Tables*. For graphical representation, data are expressed as mean ± SEM, with scatter plots overlaid to show individual data points.

## Results

### Expression of multiple microglia-signaling genes were altered within hippocampus by selective breeding for novelty responses

Our first goal was to confirm and extend findings from the meta-analysis of hippocampal gene expression in the bHR/bLR bred lines relating to microglia signaling (Birt et al., 2020). qPCR was used to measure expression levels of multiple genes within the classical complement cascade, including cascade components *C1q* (*C1qA*: NM_001008515; *C1qC*: NM_001008524) and *C3* (NM_016994), and the receptor molecule *cd11b* (NM_012711). In the brain, this canonical immune pathway represents a major signaling mechanism for microglia, reflecting inflammatory and macrophagic responses, as well as broad functions related to neurodevelopment and plasticity (Stevens et al., 2007a; Stephan et al., 2012). Compared to bHRs, hippocampus samples from bLRs had higher expression levels of multiple genes within the classical complement cascade (Figure 2, Table S3). These data confirm previously described bLR elevations in the expression of the initiator of the complement cascade, *C1qA* (*p* < 0.0001) and *C1qC* (*p* = 0.0003) (Birt et al., 2020), and extend this pattern to include the downstream effector *C3* (*p* = 0.0372), as well as the complement receptor molecule *cd11b* (*p* = 0.0021). These data implicate microglia as a source of variation that may underlie divergent behavioral traits or temperament.

**Figure 2.**
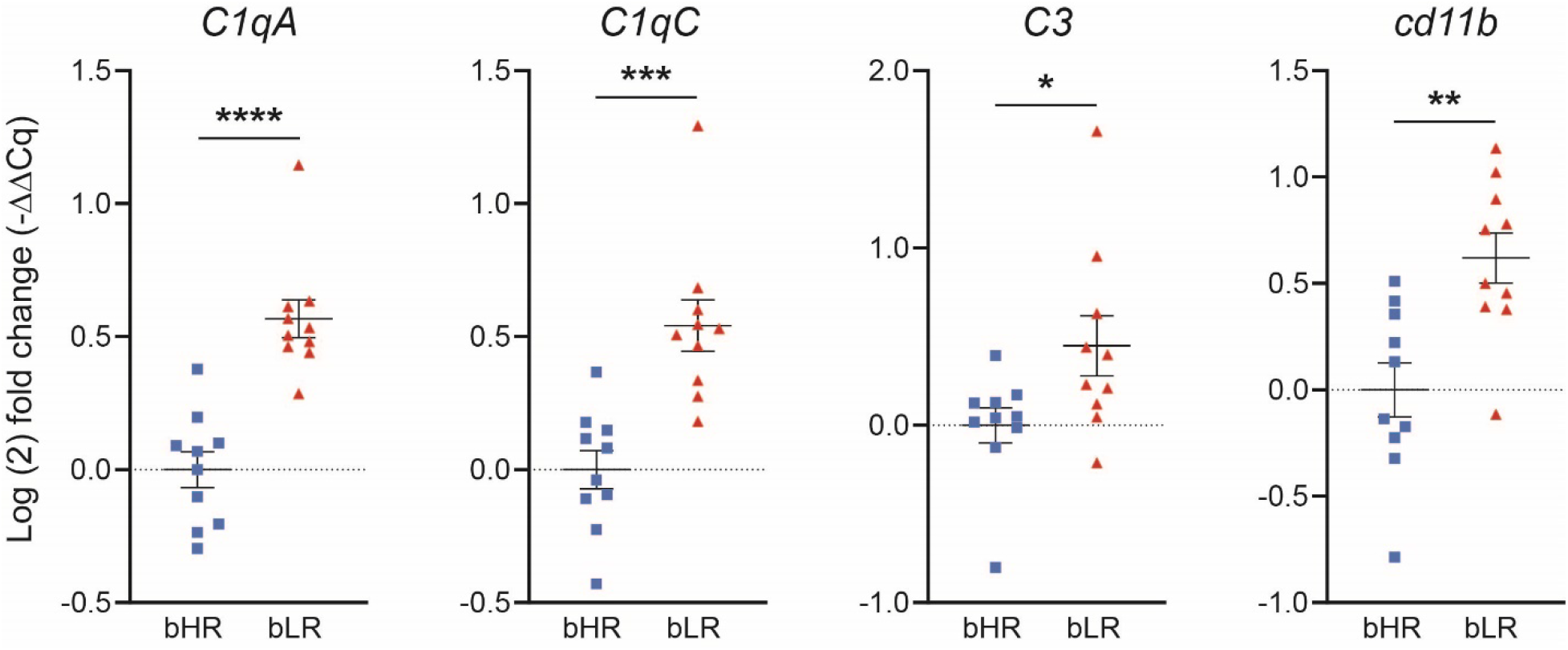
Differential patterns of microglia-related gene expression in hippocampus following selective breeding. qPCR analyses found elevations in gene expression (Log(2) fold change) in hippocampus of several genes within the classical complement cascade in bLRs compared to bHRs. *Gapdh* was used as reference gene and was not significantly different between lines (*Supplemental Materials*). Welch’s t-test compared lines, \**p* < 0.05, \*\**p* < 0.005, \*\*\**p* < 0.0005, \*\*\*\**p* < 0.0001.

### Minocycline administration improved measures of emotional and social behaviors in animals with an inhibited temperament

To test whether microglial activation plays a role in the bLR phenotype, we used minocycline administration in the drinking water to broadly inhibit microglia (Hinwood et al., 2012, 2013). Minocycline is a tetracycline antibiotic that readily crosses the blood brain barrier, and with peripheral administration, has been shown to inhibit microglia activation in the brain (Yrjänheikki et al., 1998; Yrjanheikki et al., 1999; Tikka et al., 2001; Hinwood et al., 2012, 2013; Wang et al., 2018). Body weight and fluid intake were monitored throughout the experiment and indicate systemic doses in the range of 66 – 84 mg/kg (mean = 75 mg/kg; Figure S2). Following minocycline administration, we observed specific shifts in the bLR behavioral phenotype (see Table S4 for full statistics). Although minocycline did not alter anxiety-like behavior in either the novelty-suppressed feeding test or the open field apparatus (Figure 3A, B, all *p* > 0.05), minocycline significantly increased social exploration during the social interaction test (Figure 3C; *p* = 0.023). This treatment effect was specific to the social stimulus, as there was no effect on exploration of the empty stimulus cage (*p* = 0.726), indicating minocycline impacted active social motivation in bLRs. In the forced swim test, control bLRs had high immobility scores (Figure 3D), consistent with their depressive-like phenotype. Minocycline reduced immobility scores (*p* = 0.012), which was associated with an increase specifically in swimming scores (*p* = 0.011). Finally, minocycline increased the preference to consume sucrose over water in the sucrose preference test (Figure 3E, *p* = 0.042), without altering the total volume of liquid consumed (not shown, *p* = 0.621). In summary, inhibition of microglia with minocycline treatment shifted multiple aspects of the bLR emotional phenotype.

**Figure 3.**
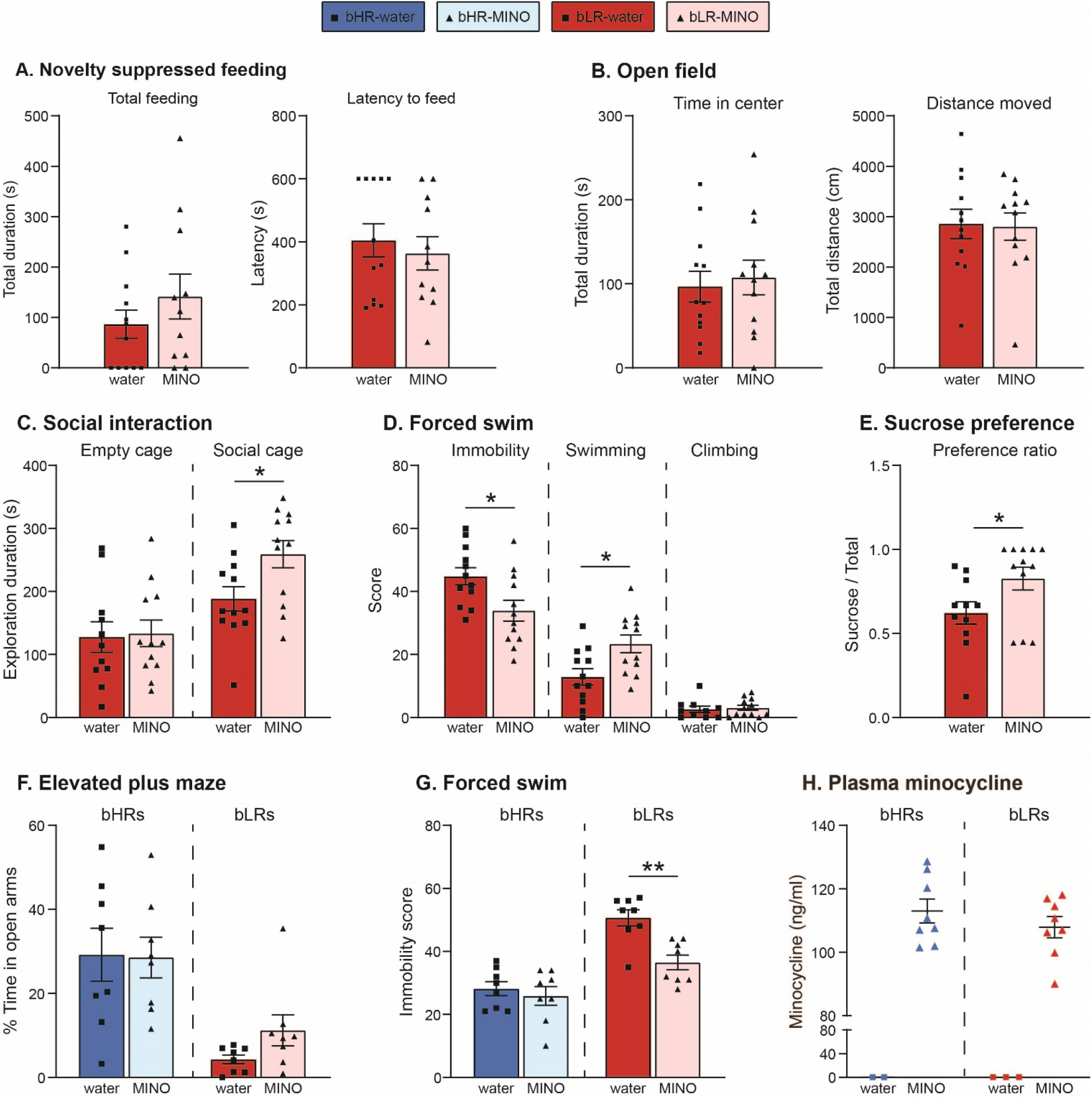
Minocycline administration altered multiple aspects of bLR phenotype. bLRs were assessed in several behavioral measures following minocycline administration (**A – E**). There was no effect of minocycline on novelty suppressed feeding task (**A**) or movement measures in the open field test (**B**) in bLRs. (**C**) In the social preference test, however, minocycline-treated bLRs spent more time exploring the cage containing the social stimulus compared to control bLRs. (**D**) In the forced swim test, minocycline administration decreased immobility scores and increased swimming scores compared to control bLRs. (**E**) In the sucrose preference test, minocycline increased the preference ratio for sucrose solution. Effects of minocycline administration were also compared between bHRs and bLRs (**F – H**). Anxiety measured in the elevated plus maze (**F**) showed that bLRs spent less time in the open arms overall, but there was no effect of minocycline treatment in either line (α reflects main effect breeding line). (**G**) In the forced swim test, there was an interaction between breeding line and treatment, such that minocycline reduced immobility scores in bLRs (** *p* = 0.001, bLR-MINO vs. bLR-water) but not in bHRs. (**H**) Plasma minocycline measurements indicated comparable levels of circulating minocycline at the end of the experiment between bHRs and bLRs. Independent *t*-tests, \**p* < 0.05, \*\**p* < 0.005 water vs. MINO treatment groups. α denotes main effect of breeding line in 2-way ANOVA.

### The behavioral effects of minocycline varied according to underlying temperament

To determine whether the observed effects of minocycline were specific to the underlying temperament of the animal, we directly compared behavior of bHRs and bLRs following a similar regimen of minocycline administration. Body weight and fluid intake were monitored throughout this experiment and found estimated comparable dose ranges in the range of 66 – 84 mg/kg (mean = 75 mg/kg; Figure S3). Consistent with line differences in anxiety-like behavior, bHRs spent significantly more time in the open arms of the elevated plus maze compared to bLRs (Figure 3F; *p* < 0.001). Minocycline had no effect on this behavior in either line (Main effect of treatment, *p* = 0.514). In the forced swim test (Figure 3G), bLRs had higher immobility scores overall compared to bHRs (*p* < 0.001), but there was also a significant interaction between treatment and line (*p* = 0.027). Simple effects analysis revealed that minocycline reduced immobility in bLRs (*p* = 0.0012) but had no effect in bHRs (*p* = 0.555), confirming the anti-depressant effect in bLRs, but no change in bHRs. Plasma minocycline levels measured in a sub-set of these subjects showed that bHRs and bLRs had comparable levels of circulating minocycline at the end of the experiment (Figure 3H, *p* > .05), indicating that the observed differences in the efficacy of minocycline did not reflect significant differences in dosing. Taken together, these behavioral studies support a role for microglia activity in shaping emotional behaviors and indicate minocycline can have differential effects according to the temperamental phenotype of the animal.

### Estimates for total hippocampal microglia number were similar in bHR and bLR lines

As part of our overall hypothesis for a role of microglia in emotional temperament, we predicted that extreme differences in inborn temperament were associated with underlying variation in microglia tone, either in the total number of microglia cells, or in their activation status. To address this aspect of our hypothesis, we generated a detailed, anatomical comparison of hippocampal microglia populations in bHRs and bLRs using immunohistochemical labeling of microglia cells. First, we calculated optical density measurements for Iba1 staining and found a trend for increased Iba1 staining density in bLRs compared to bHRs (Figure 4A; *t*_15_ = 2.038, *p* = .056). As a direct measure of the number of microglia cells across the hippocampus, we then used non-biased stereological sampling techniques to estimate total microglia number within each sub-region. For this data set, the effects of breeding line did not vary by sub-region analyzed (all line *X* region effects *p* > 0.05; Table S6); data were therefore combined across sub-regions for each subject prior to final analyses and presentation. The estimates for total number of microglia were similar between bHRs and bLRs, although there was a trend for higher totals in bLRs (Figure 4B; *p* = .085). When estimates were broken down by morphological category, for both lines, most microglia were classified as ramified (bHRs: 91.34 ± 1.44%; bLRs: 89.43 ± 1.63%). Again, estimated total numbers for each category were not significantly different between bHRs and bLRs (Figure 4B; all *p* > .05). Importantly, these direct counts of microglia cells revealed that the observed differences in microglia gene expression do not reflect differences in total microglia numbers or shifts in the bLR microglia population toward the reactive/amoeboid states.

**Figure 4.**
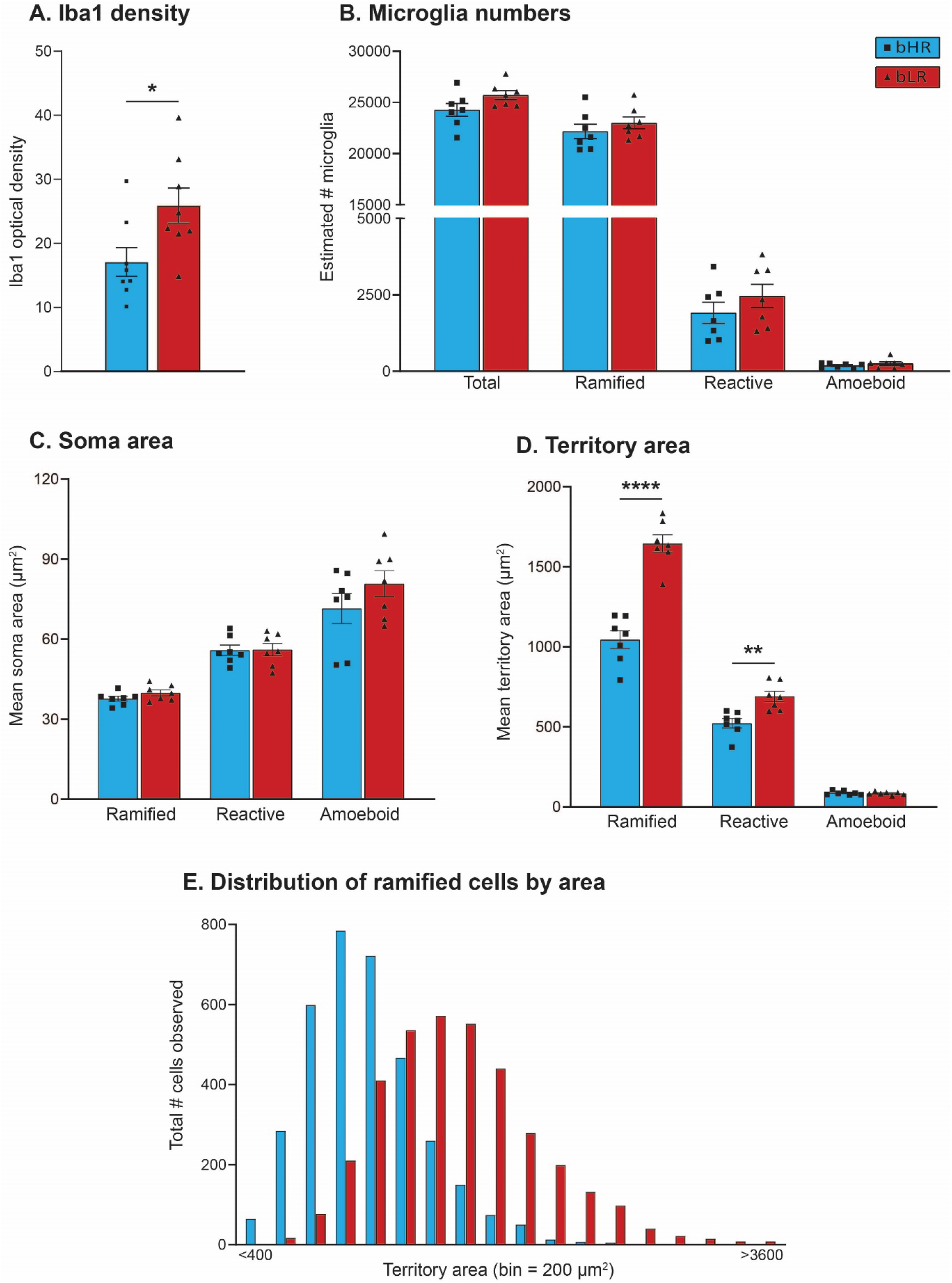
Comparison of microglia density, numbers, and area in hippocampus of bHRs and bLRs. (**A**) Optical density measurements revealed a trend (*p* = 0.056) for higher density of Iba1 staining in hippocampus of bLRs compared to bHRs (**B**) Stereological estimates of total microglia numbers were similar between bHRs and bLRs, regardless of cell type. Soma area (**C**) and territory area (**D**) were also measured for each cell counted in the stereological sampling, and the average area measurements from all cells for each subject are shown, according to morphological category. Although estimates of soma area were not different between the lines, estimates for territory area were larger in bLRs compared to bHRs for ramified and reactive cell types. (**E**) A frequency distribution of territory area for all ramified microglia measured was plotted and illustrates the right-ward shift in the overall territory size of ramified microglia from bLRs compared to bHRs. Independent *t*-tests compared bHRs v. bLRs, * *p* < .05 ** *p* < .005; **** *p* < 0.0001.

### Temperamental differences were associated with shifts in the territory area of microglia cells within hippocampus

As an additional measurement of microglia, soma and territory area estimates were generated for every cell counted within the stereological sampling procedure. This analysis generated quantitative area estimates for a range 480 – 624 total cells per subject (average cells measured per subject ± SEM: bHRs = 536 ± 14; bLRs = 567 ± 11; see Table S7 for full breakdown by region and cell type). Average soma and territory areas were calculated for each subject separately for each cell type (*Ramified, Reactive, Amoeboid*). For this data set, the effects of breeding line did not vary according to sub-region (Table S8), and area values were again collapsed across sub-regions for final analysis. In general, area estimates were consistent with criteria used for morphological categories, with increasing soma area and decreasing territory area from ramified, to reactive, to amoeboid cells (Figure 4C, D). Although there were no significant differences between bHRs and bLRs in soma area for any of the microglia types (Figure 4C; all *p* > .05), territory areas were larger in bLRs compared to bHRs for both ramified (*p* < 0.0001) and reactive (*p* = 0.0023) microglia (Figure 4D). To further illustrate differences in the size of ramified microglia across the entire population of cells measured, we generated frequency distributions of the total number of ramified microglia across the range of territory areas (binned by 200 μm^2^ increments; Figure 4E). These plots demonstrate that although territory area fell along a normal distribution in both lines, there was a right-ward shift in the bLR population: the distribution from bLRs had a larger mode and larger maximum measurements compared to the population from bHRs, again indicating that overall, ramified cells from bLRs had larger territories than those from bHRs.

### Detailed morphological analysis indicated hyper-ramification of microglia in bLRs compared to bHRs

Intriguingly, the larger area measurements of ramified cells in bLRs are consistent with elongation of microglia processes that occurs when cells undergo hyper-ramification. This unique morphology has been observed when animals are exposed to certain forms of stress or environmental challenges and is thought to reflect an intermediate stage of the microglia response (Hinwood et al., 2012, 2013; Walker et al., 2014). To directly assess the possibility of hyper-ramification in bLRs, we performed a more detailed morphological analysis of 3-D cell reconstructions generated for a sub-set of ramified microglia from bHRs and bLRs. For this data set, breeding line effects were not homogenous across sub-region (Table S8), and the data were therefore analyzed and presented separately for CA1, CA3, and DG. Figure 5A and B show high magnification photomicrographs of typical ramified cells from each line, along with their associated reconstructions generated using Neurolucida (Figure 5C, D). Overall, the morphological analyses were consistent with the process area estimates described above, and importantly, provide more detailed information regarding microglia complexity between breeding lines. Although the total number of primary processes per cell was similar between the lines (Figure 5E; all *p* > .05), the average length of these processes was longer in bLRs compared to bHRs (Figure 5F, CA1, *p* = 0.001; CA3, *p* < 0.001). The number of branch points per cell, reflecting process complexity, was also higher in bLRs compared to bHRs (Figure 5G; all sub-regions *p* < 0.001). As a result of the longer and more complicated processes in bLRs, the area of the territory encompassed by the entire cell (estimated using convex hull analysis) was larger in bLRs compared bHRs (Figure 5H), specifically within CA1 (*p* < 0.001) and CA3 (*p* < 0.001). Soma area from these reconstructions revealed a significantly larger soma area in bLRs compared to bHRs within CA1 and CA3 (Table S9; CA1, *p* = 0.008; CA3, *p* = 0.003). In summary, microglia from bLRs had longer processes, with more complicated branching patterns, and larger overall territories, an overall pattern that is consistent with a hyper-ramified state.

**Figure 5.**
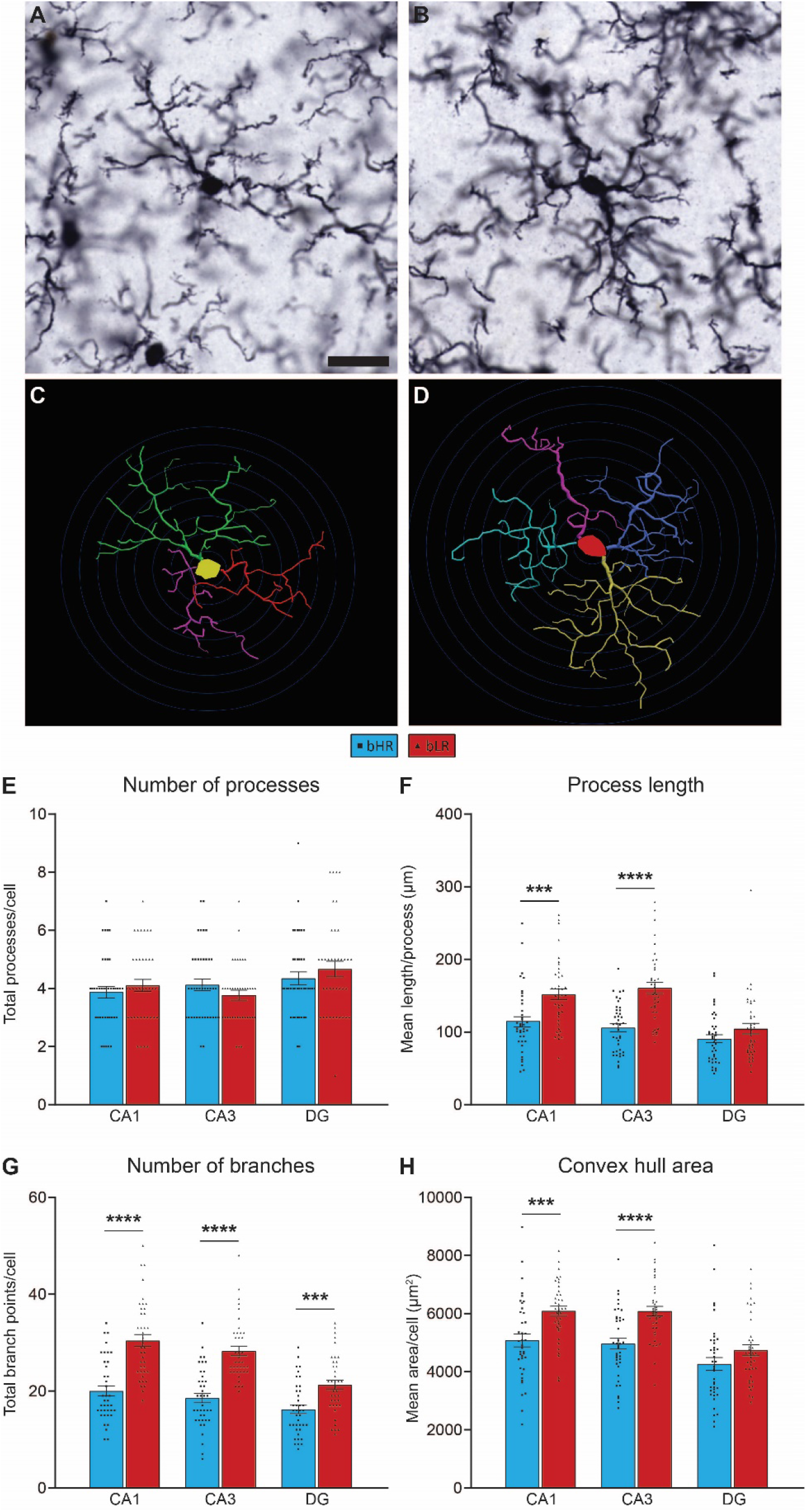
Detailed morphological analyses of ramified microglia in bHRs and bLRs. High-magnification photomicrographs (63-x, oil) of typical ramified microglia cells from bHR (**A**) and bLR (**B**), with Neurolucida reconstructions for each cell shown in (**C**) and (**D**), respectively. Scale bar = 20 μm; Concentric circles applied at 5 μm increments in **C** and **D** for length comparisons. Analyses revealed that although microglia from each line had similar totals of primary branches (**E**), bLR microglia had longer branches in CA1 and CA3 (**F**), and more branch points across all regions (**G**), compared to microglia from bHRs. (**H**) Convex hull calculations, which reflect the overall territory area of the cell, were larger in bLRs in CA1 and CA3. Independent *t*-tests compared bHRs v. bLRs, *** *p* < .0005; **** *p* < 0.0001.

## Discussion

The current set of experiments used a selective breeding model to examine the role microglia play the hippocampus of animals genetically selected for contrasting temperaments. The bHR and bLR lines have been bred for differences in their response to novelty and reflect a range of divergent emotional responses, including anxiety- and depression-like behaviors (Turner et al., 2017). After identifying elevated expression levels of several microglia-related genes from hippocampus of bLRs, we hypothesized that microglia play a role in shaping their uniquely inhibited phenotype. We found that the microglia blocker minocycline improved several emotional measures in bLRs, with no observable effects in bHRs. To determine precisely how the microglia populations differ between the bred lines, we performed detailed anatomical comparisons of immunohistochemically labeled microglia. These analyses found that, although bHRs and bLRs had comparable total numbers of microglia within hippocampus, cells from bLRs were characterized by a hyper-ramified morphology, reflective of an intermediate stage of microglia activation. This study provides evidence linking variation in inborn temperament with differences in the baseline morphological status of microglia. Our findings highlight the importance of considering microglia as key players in understanding differences enduring differences in emotional responsivity.

### Animals with divergent temperaments express differential levels of microglia-signaling genes within hippocampus

Over the fifteen years of research on bHR and bLR lines, multiple molecular pathways have been identified underlying these unique phenotypes (Ballaz et al., 2007, 2008; Perez et al., 2009; Turner et al., 2011, 2017). A recent meta-analysis compared gene expression profiles collected from 8 transcriptional datasets, spanning 43 generations of selective breeding, and found a broad range of hippocampal genes associated with breeding (Birt et al., 2020). Collectively, these studies confirm a role for the hippocampus in the behavioral phenotypes, as well as highlight novel candidate pathways to explore. Of these potential candidates, microglia-related genes were found to be upregulated in bLRs compared to bHRs, a pattern that complements the emerging theories of microglia mechanisms of mood disorders (Yirmiya et al., 2015). One of the top microglia genes identified by this analysis, *C1q,* plays a critical role in microglia signaling as the initiator of the classical complement cascade (Stevens et al., 2007b; Stephan et al., 2012). The meta-analysis also showed that a genetic variant near *C1qa* (on chromosome 5) segregates bHR and bLR lines, providing evidence for genetic determinants of the differential expression profiles (Birt et al., 2020).

The current qPCR analyses of hippocampus from bHRs and bLRs confirms the differential expression of *C1q* observed in previous transcriptional datasets and further extends the set of genes to include additional molecules within the classical complement pathway. In addition to *C1q,* the *C3* ligand and a component of its receptor *cd11b* (CR3) was also upregulated in bLRs. It should be noted that across expression profiles done to-date (Birt et al., 2020; and unpublished data), there has been no evidence of increased expression of traditional neuroinflammatory markers, including pro-inflammatory cytokines. This finding is consistent with reports that bHR/bLR lines have similar levels of plasma cytokines (Glover et al., 2021) and indicate the functional significance of bHR/bLR differences in microglia gene expression may reflect non-pathological, nonimmune functions. Microglia signaling is known to play a role in many aspects of healthy brain function, and the complement cascade pathway in particular has been linked to synaptic pruning in development and plasticity throughout life (Stevens et al., 2007b; Schafer et al., 2012; Stephan et al., 2012). Determining the specific role for the complement pathway in regulating emotional phenotypes in our selective breeding model is a compelling avenue for future studies.

### Minocycline treatment alters key aspects of the inhibited phenotype of bLRs

Research into the use of minocycline for treatment of affective disorders is expanding, and a meta-analysis of random controlled clinical trials of minocycline suggest good therapeutic potential in patients with major or bipolar types of depression (Rosenblat and McIntyre, 2016, 2018). The clinical trials are supported by many rodent studies that report anti-depressant effects of minocycline (Henry et al., 2008; Molina-Hernández et al., 2008; Pae et al., 2008; O’Connor et al., 2009; Arakawa et al., 2012; Hinwood et al., 2012, 2013; Zhu et al., 2014; Kreisel et al., 2014; Zheng et al., 2015; Majidi et al., 2016; Tong et al., 2017; Wang et al., 2018; Zhang et al., 2018, 2019b, 2019a). Although some anti-depressant effects have been observed in naïve animals (Molina-Hernández et al., 2008), most studies report behavioral changes specifically following a pro-depressant challenge (e.g., chronic stress, immune stimulation). The fact that minocycline typically has no effects in control animals suggests that outbred strains may not be sensitive to the effects of minocycline at baseline conditions. However, the natural variation that exists within outbred strains leaves the possibility that minocycline may be efficacious in certain sub-populations, but those effects are not detected when averaged across the whole group.

In support of this interpretation, we found that minocycline had good efficacy in emotionally vulnerable bLRs, even in the absence of any overt stress or challenge. Minocycline improved bLR behavior in multiple depressive-like measures, whereas in bHRs, minocycline had no effect in the forced swim test, one of the most commonly used tests for anti-depressant efficacy (Slattery and Cryan, 2012). Similar to the results described here, Schmidtner et al (2019) also showed that minocycline reduces depressive-like behaviors in rats bred for high anxiety behavior (HAB), but not in the non-anxious line. Indeed, the current behavioral effects line up well with those described in the HAB rats, finding consistent effects of minocycline in both the forced swim and social exploration tests, and no changes in anxiety measures. In addition, our study reported minocycline increased sucrose preference in bLRs, expanding the effects to include anhedonia measures of depression. Together, the results from these selective breeding models highlight the importance for understanding how inborn traits may shape the expression of emotional behaviors and sensitivity to therapeutics.

Minocycline has well-established functions of suppressing microglia proliferation and signaling (Yrjänheikki et al., 1998; Yrjanheikki et al., 1999; Tikka et al., 2001). As a result, much discussion around the effects of the drug center around its role in inhibiting microglia function. The tight link between microglia and depressive behavior observed in various animal models (Pae et al., 2008), and the consistent shift in both behavior and microglia following minocycline administration (Hinwood et al., 2012; Wang et al., 2018; Zhang et al., 2019b), suggest a functional relationship. It is important to note, however, that minocycline may have broad effects, with direct and indirect consequences extending beyond microglia function. In addition to its anti-inflammatory effects, potential anti-depressant mechanisms of minocycline include regulation of neurogenesis, antioxidation, apoptosis, and excitatory toxicity processes (Yong et al., 2004; Pae et al., 2008). Minocycline may even be acting outside of the brain by changing the composition of the gut microbiome (Schmidtner et al., 2019), although a recent study in the bHR/bLR lines suggests that these temperaments are not associated with baseline differences in their microbiota (Glover et al., 2021). The precise mechanism through which minocycline alters emotional behavior will need to be explored further in this model and may involve dynamic interactions among multiple pathways.

### Morphological status of microglia cells is shifted in animals with divergent temperaments

Microglia are dynamic players in the brain, and their function can be reflected by alterations in total numbers, as well as functional state. When at rest, microglia cells maintain a ramified structure, with long motile processes actively surveying their local territory (Nimmerjahn et al., 2005; Kettenmann et al., 2011, 2013). Upon stimulation, microglia morphology shifts along a continuum, progressing through a reactive phase, with shortening and thickening of processes, to the fully activated amoeboid (macrophagic) phase, with enlarged soma and full retraction of processes (Streit et al., 1999; Stence et al., 2001; Walker et al., 2014). Descriptions of microglia “activation” typically refer to these reactive/amoeboid morphologies, yet a growing body of research recognizes additional morphological markers within the microglia response (Walker et al., 2014). In particular, the “hyper-ramified” morphology, characterized by an elongation of processes and increase in branch complexity, is thought to reflect an early stage of activation, or perhaps occurs in response to relatively less intense (non-pathological) stimulation (Hinwood et al., 2012, 2013; Walker et al., 2014). A major goal of the current study was to perform detailed comparisons of bHR/bLR, both in terms of microglia numbers and morphological characteristics, to determine how selective breeding may have shifted the tone of their microglia populations.

Interestingly, our anatomical studies revealed that microglia numbers were not affected by selective breeding. In fact, bHRs and bLRs had similar total numbers of microglia, as well as similar numbers of cells categorized in the reactive or amoeboid states. This result suggests that the elevated expression of microglia-related genes observed in bLR hippocampus does not reflect underlying differences in microglia number or shifts in their cells toward fully activated morphologies. Further detailed analyses, however, identified significant differences in microglia morphology: ramified microglia from bLRs had longer processes, with more complicated branch patterns, than microglia from bHRs. Microglia from bLRs also had larger overall territories, a difference detected across large numbers of cells measured during stereological sampling, as well as from the detailed reconstructions of a sub-set of ramified microglia. The morphology observed in bLR microglia is consistent with what has been described for hyper-ramification and suggests ramified cells in this line are not in a truly quiescent phase.

A growing number of studies has now reported evidence of hyper-ramified microglia following environmental challenges. Manipulations that provoke hyperramification include various forms of chronic stress, including restraint, inescapable swim, foot shock, and prenatal dexamethasone exposure (Hinwood et al., 2012, 2013; Walker et al., 2014; Hellwig et al., 2016; Caetano et al., 2017; Ganguly et al., 2018; Duarte et al., 2019; Smith et al., 2019), an interesting common thread given the importance of stress as a factor for mood disorders (Mcewen and Akil, 2020). In one particularly relevant study, wild-type mice exposed to repeated forced swim display increased despair behavior, concomitant with increased hyper-ramification within dentate gyrus; both the behavioral and microglia effects are abrogated by treatment with the antidepressant venlafaxine (Hellwig et al., 2016). Using a model of chronic restraint stress, Hinwood and colleagues (2012, 2013) describe a pattern of hyper-ramification within prefrontal cortex that is reversed by minocycline treatment, suggesting a potential mechanism relevant to the current study. These previous studies provide compelling evidence that hyper-ramification signifies a unique and critical feature within the range of microglia responses to stress, and furthermore, is causally linked to changes in behavior.

Our results add to that growing body of knowledge in an important way-to our knowledge, the current data provide the first evidence associating hyper-ramification status *at baseline* to emotional vulnerability. Indeed, in our anatomical studies, there was no overt experimental stress or manipulation, and we did not see evidence for later stages of microglia activation, suggesting that we captured the steady-state of microglia in these selectively-bred lines. There is limited data on the range of microglia territory sizes expected across large populations of cells, and it can be difficult to compare across different methods of measurement. Although ramification was not examined directly, a study in rat prefrontal cortex described surprisingly large variation in the measured territory area of ramified microglia (Kongsui et al., 2014), consistent with the range of areas observed here. Our data suggest that within the range of microglia ramification, the extreme ends of that spectrum are associated with divergent emotional temperaments. Moreover, the microglia profile associated with the inherently inhibited temperament at baseline closely matches the microglia profiles described following stress, highlighting a compelling convergence between genetic and environmental models of emotional regulation.

What are the consequences of hyper-ramification on microglia physiology and function? Hyper-ramification has not been associated with markers of neuroinflammation or neurotoxicity (Sugama et al., 2007; Hinwood et al., 2012, 2013), suggesting a link with non-diseased conditions. This morphology may reflect a transitional step intermediate to full activation phases, or it may represent its own distinct endpoint. One possibility is that hyper-ramified microglia are in a “primed” state and would have heightened responses to future perturbations. In a mouse PTSD model, increased hyper-ramification within hippocampus was associated with a decrease in spine density in the same brain region (Smith et al., 2019), fitting with an interpretation of over-active macrophagic pruning responses by hyper-ramified microglia. In fact, hyper-ramification has been shown to be associated with increased expression of C1q (Smith et al., 2019) and CR3 (Streit et al., 1999; Sugama et al., 2007; Graeber, 2010), both known to serve synaptic pruning functions via their roles in the classical complement cascade (Stevens et al., 2007b; Schafer et al., 2012; Shi et al., 2015; Hong et al., 2016). Alternatively, it is possible that hyper-ramified microglia are arrested in this phase, and subsequent physiological responses may be impaired. In support of this possibility is the fact that in the aged brain, microglia can develop extreme hyper-ramification, and as a result, can become tangled and dysfunctional (Streit and Sparks, 1997). Although we did not observe evidence for microglia tangles in our lines, it is interesting to consider the possibility that inborn differences in microglia morphology may reflect an accelerated aging process. Future studies will be needed to determine the significance of microglia hyper-ramification in bHRs and bLRs and on emotional behavior in general. Given the dynamics and complexity of microglia, the consequences of hyper-ramification will likely vary according to context of the model (type of manipulation), developmental window (development v. adult), and timing of response (baseline, acute, chronic).

### Concluding remarks

Individual differences in temperament represent stable traits that shape patterns and style of reactivity to the environment in an ongoing manner as well as under significant physical and psychosocial challenges. Understanding the genetic and neurobiological mechanisms that underlie differences in temperament is essential to uncovering the biology of mood and other affective disorders. Through our selective breeding rat model, we have described an inhibited temperament that is genetically based, highly stable, and tracks with behavioral vulnerability across multiple measures. The current study adds to this knowledge by linking a unique, hyper-ramified microglia profile with that inhibited temperament. Our work shows that the baseline morphology and status of microglia has a genetic basis, that differences in microglia morphology are not only modulated by environmental stressors, but may well represent an endophenotype that biases emotional responsiveness and coping style in an ongoing manner.

