## Supplemental Materials for "Differences in microglia morphological profiles reflect divergent emotional temperaments: Insights from a selective breeding model"

### Supplemental Materials and Methods

*bHR/bLR breeding colonies*

The bHR and bLR lines were initially generated from commercially purchased Sprague-Dawlye rats (Charles River, Inc). Adult rats were screened for their exploratory response to novelty by placing them in a novel cage and the top and bottom 20% of locomotor responders were used to breed bHR and bLR lines, respectively. Locomotion was measured by individually placing an adult rat into a novel, standard housing cage, with no bedding and a grated floor. Locomotion in the novel cage was recorded every 5 minutes for 1 hour by photocells positioned along the sides of the cage connected to a computer. Over the course of subsequent generations, we have maintained 12 separate breeding families for each line, selectively breeding rats with locomotor scores (total beam breaks) closest to their respective phenotypic score and limiting inbreeding (Stead et al., 2006). In current breeding generations, average total beam breaks for bHRs and bLRs were 1800 and 150, respectively.

All litters were culled to 6-12 pups per litter (sex ratio approximately 1:1) by postnatal day 3. Rats were weaned on day 21, pair-housed with a rat from the same breeding line and left undisturbed until experimental use in adulthood. Rats were housed on a 12:12 light:dark schedule (lights on 6a.m., Standard Time), with *ad libitum* access to food and water, unless otherwise noted.

*qPCR experiments*

Tissue collection for qPCR experiments took place between 9:00 am and 2:00 pm. Rats were brought into a waiting room immediately adjacent to the procedure room used for collection and allowed to sit undisturbed for at least 1 hour prior to sacrifice. Rats were sacrificed via rapid decapitation without anesthesia and hippocampus was dissected on ice within 1 – 2 minutes following decapitation. Whole hippocampal samples from each hemisphere were placed into separate sterile collection tubes and immediately frozen on dry ice. Samples were stored at −80°C until further processing.

Hippocampal samples were homogenized using a motorized pestle, followed by Qiashredder columns (Qiagen # 79654). RNA was extracted using the RNeasy Mini Kit (Qiagen #74104), with an additional step of DNAse digestion (Qiagen #79254). RNA concentrations were measured via spectrophotometry, and cDNA was synthesized from 400 ng of template RNA (per 20 µl reaction) using the iScript cDNA Synthesis Kit (Biorad #1708891). Resulting cDNA was diluted (1:10) with RNAse-free water and stored in aliquots at −20°C.

Primers were either custom designed using Basic Local Alignment Search Tool (BLAST) through the National Center for Biotechnology Information (NCBI;

www.ncbi.nlm.nih.gov) or using published primer sequences (see Table S1). Primer sequences were checked via BLAST for specificity of sequence alignment and ordered through IDT. All primers were validated in-house by performing melt curves analysis on amplification specificity and generating amplification efficiency curves based on standard dilutions of stock cDNA. Amplification efficiencies for each primer pair were confirmed ≥ 90% (Figure S1).

Amplification reactions were carried out in 96-well PCR plates (Bio-Rad #HSP9601), and each well was loaded with a 20-μl reaction containing a final concentration of 250 nM forward and reverse primers, 50% SYBR® Green master mix (Bio-Rad #1708880), and 5% template cDNA. Reactions were run through 40 amplification cycles (95°C for 15 s, 60°C for 15 s and 72°C for 15 s) and minimum quantification cycle (Cq) for detection was determined. Samples were run in duplicate, and any samples that had ≥ 0.25 Cq discrepancy between replicates were removed from the analysis. Mean Cq values were then analyzed using the Livak method (Livak & Schmittgen, 2001), with *Gapdh* as the reference gene and bHRs assigned as the comparison group. ΔCq and ΔΔCq values were calculated for every sample, where $\Delta Cq = Cq_{\text{(target gene)}} - Cq_{\text{(Gapdh)}}$ and $\Delta\Delta Cq = \Delta Cq_{\text{(individual sample)}} - Cq_{\text{(mean comparison group)}}$. Data were plotted as inverse Log(2) fold change relative to bHR group (-ΔΔCq). Comparisons between bHRs and bLRs in $\Delta Cq_{\text{(Gapdh)}}$ values indicated no significant group difference in expression of the reference gene (Figure S1).

*Brain tissue fixation via transcardial perfusion*

All perfusions took place between 9:00 am and 2:00 pm. Rats were injected with an overdose of sodium pentobarbital (100 mg/kg), and upon reaching a deep level of anesthesia, were transcardially perfused with 200 ml of 0.1M phosphate-buffered saline (PBS; pH 7.4), followed by 200 ml of 4% paraformaldehyde in phosphate buffer (pH 7.2). Brains were immediately removed and post-fixed in 4% paraformaldehyde overnight (4°C) and then transferred to 30% sucrose in PBS for 48–72h (4°C). Brains were frozen in 2-methyl butane (-20°C) and stored at -80°C until sectioning.

*Iba-1 immunohistochemistry*

All tissue was processed concurrently, and master mixes of all solutions were used to reduce variability across wells. Unless otherwise stated, all rinses and incubations were performed with gentle agitation and at room temperature. Sections were first rinsed thoroughly in PBS (pH 7.4). To enhance the antigenicity of the endogenous Iba1 signal, sections were incubated in 10mM sodium citrate (pH 9.0) for 30 minutes at 80°C (Jiao et al., 1999). After several rinses in PBS, tissue sections were incubated in 0.5% $H_2O_2$ in PBS for 15 min and rinsed again in PBS. Sections were incubated in the primary antibody against Iba1 at 1:70,000 concentration (019-19741; FUJIFILM Wako

Chemicals, Richmond, VA) in PBS with 0.4% Triton X-100 for 1 h at room temperature, and then for 48 h at 4°C. Following primary incubation, sections were rinsed in PBS and then incubated for 1h in an anti-rabbit biotinylated secondary antibody (1:600; Jackson ImmunoResearch, West Grove, PA) in PBS with 0.4% Triton X-100. Sections were rinsed again in PBS, and then incubated for 1h in avidin–biotin complex (4.5μL each of A and B reagents/mL of PBS with 0.4% Triton X-100) (ABC Elite Kit; Vector Laboratories, Burlingame, CA). Sections were rinsed in PBS, followed by 0.175M sodium acetate, and were then incubated in nickel sulfate (25 mg/mL; Sigma-Aldrich, St. Louis, MO), 3,3-diaminobenzidine-HCl (0.2 mg/mL; Sigma-Aldrich), and 0.05% $H_2O_2$ in sodium acetate for 15 min. The staining reaction was stopped by rinsing sections in sodium acetate, followed by PBS. Stained sections were mounted onto electrostatically charged glass slides and allowed to air-dry overnight. Slides were then dehydrated in alcohols, cleared in xylenes, and coverslipped with Permount.

*Behavioral assays*

All behavioral tests were conducted between 9:00 am and 1:00 pm and sequential tests were separated by 24 – 72 hours. Testing was done under overhead white fluorescent lighting, unless otherwise specified. All behavioral coding was done by a researcher who was blind to the identity of the subject.

*Novelty-suppressed feeding test.* Rats were food-deprived on the night prior to testing to increase motivation to feed. Immediately prior to testing, 3 pellets of standard chow were placed into the center of a novel testing arena (33 x 60 cm), fitted with overhead video recorders. Rats were placed into the arena along the back wall and were allowed to explore the arena for 10 minutes. The latency to feed and total duration of feeding was later scored by a researcher using Observor software (Noldus).

*Social interaction in an open field.* Rats were tested in an open field apparatus (100 x 100 cm) with a black matte floor and white walls under dim light (40 lux). The testing consisted of two phases, distinguished only by the presence or absence of a social stimulus. The first phase was used to both habituate the subject to the apparatus and also provide open field exploratory data. During this phase, the subject was placed into the open field with an empty wire mesh chamber positioned along the north wall. At the end of 10 minutes, the subject was briefly removed and placed into a holding cage, while the empty mesh container was replaced with a similar container holding a single social stimulus rat. The stimulus rat was a novel, commercially purchased (Charles River Laboratories, Wilmington, MA) Sprague Dawley rat of the same age and approximate weight as the subject. The wire mesh permitted transmission of visual and auditory cues, while limiting direct physical contact. The subject was then placed back into the open field apparatus now containing the social stimulus and allowed to explore for another 10 minutes. The center-location of the subject was automatically tracked during both phases using Ethovision (Noldus). In addition, the total duration of direct exploration of the stimulus chambers was later scored by a researcher using Observor (Noldus). Direct exploration was scored when the subject: climbed on top of the stimulus cage; directly contacted the stimulus cage with its front paw(s); or directed its nose within 1 cm of any side of the stimulus cage.

*Sucrose preference test.* On the day prior to sucrose preference testing, rats were habituated to the sucrose solution by replacing both of their home cage water bottles with bottles containing 1% sucrose (in tap water) for 4 hours. That night, all rats were food and water deprived (16 hours) to increase motivation to drink. On the morning of sucrose preference testing, each rat was transferred into a clean home cage individually for a 2-hour sucrose preference test, during which time they could drink freely among two bottles: one containing water and the other containing 1% sucrose solution. Bottles were weighed immediately before the test, and then again at the end of the test. Sucrose preference ratios were calculated by dividing the volume of sucrose solution consumed by the total volume of liquid consumed.

*Forced swim test.* Rats were tested for swim behavior as their final assay using a modified Porsolt protocol (Slattery & Cryan, 2012). Rats were placed into large opaque plastic cylinders containing 35-cm deep water (25 °C ± 1 °C). On Day 1, rats were placed in the cylinder for 15 minutes; on day 2 (24 h later), rats were placed in the cylinder for 5 minutes. Water was emptied after each swim, so every rat was tested with fresh water. Overhead video recorders were used to record the tests, and videos were later scored using Observor (Noldus). Scoring was performed using a time-sampling technique, wherein the predominant behavior of each 5-second period was recorded throughout the duration of the 5-minute test, for a total of 60 samples. Active swim behaviors included swimming (horizontal movements at the surface of the water), climbing (upward-directed movements against the cylinder using forepaws), or diving (swimming directed toward the bottom of the tank). Immobility (or passive swim) behavior was defined as floating without struggle and only making minimal movements necessary to keep head above water.

*Elevated plus maze.* Rats were placed in the center of an elevated plus maze constructed of black Plexiglass (45 cm arms, elevated 70 cm from floor) illuminated with dim light (40 lux). Rats were given 5 minutes to explore the maze, during which time their center-body location was automatically tracked using an overhead video camera connected to a computer with Ethovision software (Noldus Information Technology, Leesburg, VA). The percentage of time spent in the open arms of the maze, as well as total distance moved, were calculated.

**Figure S1. Validation of primer sequences used in qPCR studies.** qPCR was used to measure expression levels of multiple genes within the classical complement cascade: *C1qA*, *C1qC*, *C3*, and *cd11b*. Primer amplification efficiency curves were generated based on standard dilutions of stock cDNA and found primer efficiencies for all primer pairs were over 93%. Furthermore, expression levels of the housekeeping gene, *GAPDH*, was similar between bHRs and bLRs ($t_{10.37} = 0.9462$, $p = 0.3656$).

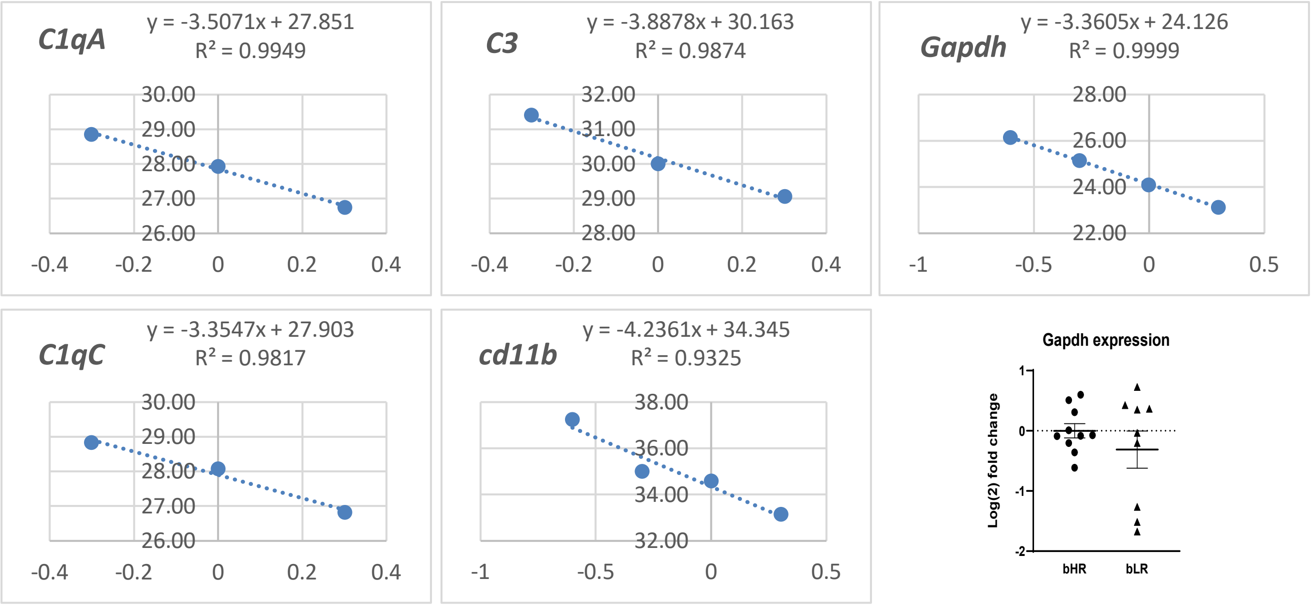

**Figure S2. Bodyweight and estimated minocycline dosage in bLR experiment.** (A) Body weight was measured periodically throughout minocycline administration and revealed comparable growth rates between bLRs given tap water or minocycline in water bottles. Using body weight and liquid consumption, average estimated minocycline dosages (mg/kg) were calculated for bLR-MINO rats and ranged from 66 – 84 mg/kg (Mean = 75 mg/kg).

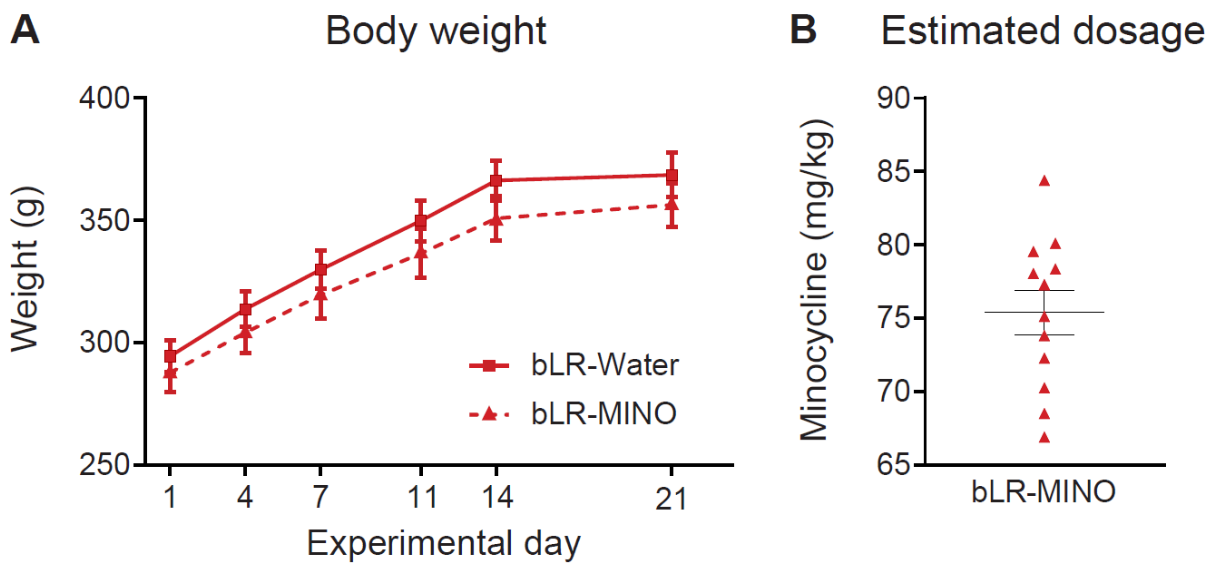

**Figure S3. Bodyweight and estimated minocycline dosage in bHR v. bLR experiment.** (A) Body weight was measured periodically throughout minocycline administration and revealed comparable faster growth rates in bHRs, which is typical in these lines. In this experiment, minocycline administration was associated with body weight in bLRs. Regardless, average estimated minocycline dosages (body weight/consumption, mg/kg) were similar in bHRs and bLRs and ranged from 68 – 98 mg/kg.

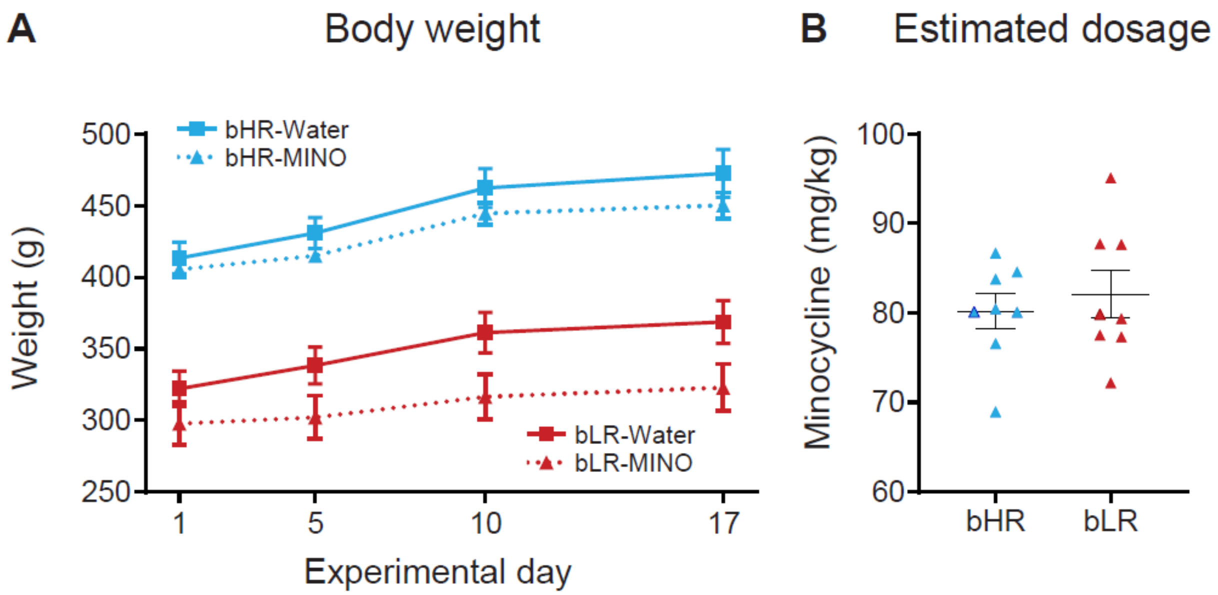

**Table S1. Primer sequences for qPCR confirmation of microglia-related genes**

| Gene | Accession ID | | Primer sequence |
|---|---|---|---|
| Complement C1q A chain (C1qa) | NM_001008515 | *F* | ACAAGGTCCTCACCAACCAG |
| | | *R* | CGTTGCAATTGAAGCACAGT |
| Complement C1q C chain (C1qc) | NM_001008524 | *F* | ACTTCGTCCACCACACATCC |
| | | *R* | ACCATGCCGTTGTAGTCGTT |
| Complement C3 (C3) | NM_016994 | *F* | TATTCCAGACACAAACGACCTG |
| | | *R* | GAACTGGTGGACTTTGAAGGAC |
| Integrin, alpha M (cd11b; Itgam; CR3) | NM_012711 | *F* | GGGAAACGCCTTCCACAAAC |
| | | *R* | GTACTTCCTGTCTGGGTGCC |
| Gapdh | NM_017008 | *F* | GTTTGTGATGGGTGTGAACC |
| | | *R* | TCTTCTGAGTGGCAGTGATG |

### Table S2. Sampling parameters used for stereological estimates of microglia

| Stereological variable | Value | |
|---|---|---|
| Number sections/subject | 4 | |
| Section sampling fraction | 1/6 | |
| Area of sampling fractions | (100 μm X 100 μm)/(250 μm X 250 μm) | |
| Height of disector | 12 μm | |
| Guard zone | 1.0 μm | |
| | **bHRs** | **bLRs** |
| Average measured thickness of section | 14.333 ± 0.087 | 14.324 ± 0.119 |
| Observed coefficient of error (Gundersen, *m* = 1) | 0.078 ± 0.001 | 0.077 ± 0.001 |

### Table S3. Statistics for qPCR studies

| Welch's *t*-test statistics: bHRs v. bLRs | *t* (df) | Significance |
|---|---|---|
| *C1qA* | 5.806 (17.92) | $p < 0.0001$ |
| *C1qC* | 4.506 (16.63) | $p = 0.0003$ |
| *C3* | 2.293 (14.52) | $p = 0.0372$ |
| *cd11b* | 3.588 (17.89) | $p = 0.0021$ |

### Table S4. Statistics for determining minocycline effects in bLRs

| **Independent *t*-test statistics: minocycline v. water** | ***t* (df)** | **Significance** |
|---|---|---|
| **B. Novelty suppressed feeding** | | |
| Feeding duration | 1.059 (21) | $p = 0.301$ |
| Latency to feed | 0.546 (21) | $p = 0.591$ |
| **C. Open field** | | |
| Time spent in center | 0.387 (22) | $p = 0.703$ |
| Total distance moved | 0.134 (22) | $p = 0.894$ |
| **D. Social exploration** | | |
| Empty cage | 0.355 (22) | $p = 0.726$ |
| Social cage | 2.454 (22) | **$p = 0.023$** |
| **E. Forced swim** | | |
| Immobility | 2.754 (22) | **$p = 0.012$** |
| Swimming | 2.775 (22) | **$p = 0.011$** |
| Climbing | 0.779 (22) | $p = 0.444$ |
| **F. Sucrose preference** | | |
| Preference ratio | 2.155 (21) | **$p = 0.042$** |
| Total consumption | 0.502 (21) | $p = 0.621$ |

### Table S5. Statistics for determining minocycline effects in bHRs and bLRs

| | Mixed ANOVAs (line x sub-region) | | | | Simple main effects | |
|---|---|---|---|---|---|---|
| Effect | *F* (df) | Significance | Partial Eta Squared | | *t* (df) | Significance |
| **Elevated plus maze** | | | | | | |
| *Percent time in open arms* | | | | | | |
| Line | 2.465 (1,12) | *p* = 0 .142 | 0.170 | | | |
| Treatment | 37.662 (2,11) | ***p* < 0.001** | 0.873 | | | |
| Line X Treatment | 0.192 (2,11) | *p* = 0.828 | 0.034 | | | |
| *Total distance traveled* | | | | | | |
| Line | 216.822 (1,28) | ***p* < 0.001** | 0.886 | | | |
| Treatment | 0.033 (1,28) | *p* = 0.857 | 0.001 | | | |
| Line X Treatment | 1.546 (1,28) | *p* = 0.224 | 0.052 | | | |
| **Forced swim** | | | | | | |
| *Immobile/Passive floating* | | | | | | |
| Line | 42.155 (1,28) | ***p* < 0.001** | 0.601 | *bHRs* | 0.605 (14) | *p* = 0.555 |
| Treatment | 10.302 (1,28) | ***p* = 0.003** | 0.269 | *bLRs* | 4.048 (14) | ***p* = 0.001** |
| Line X Treatment | 5.418 (1,28) | ***p* = 0.027** | 0.162 | | | |
| *Swimming* | | | | | | |
| Line | 18.132 (1,28) | ***p* < 0.001** | 0.393 | *bHRs* | 0.669 (14) | *p* = 0.515 |
| Treatment | 8.821 (1,28) | ***p* = 0.006** | 0.240 | *bLRs* | 8.611 (14) | ***p* < 0.001** |
| Line X Treatment | 17.528 (1,28) | ***p* < 0.001** | 0.385 | | | |
| *Climbing* | | | | | | |
| Line | 0.863 (1,12) | *p* = 0.606 | 0.633 | | | |
| Treatment | 0.556 (2,11) | *p* = 0.688 | 0.526 | | | |
| Line X Treatment | 0.067 (2,11) | *p* = 0.820 | 0.032 | | | |

### Table S6. Statistics from cell number analyses

| Mixed ANOVAs (Line x Sub-region) | | | | Independent t-tests (bHR v. bLR) | |
|---|---|---|---|---|---|
| **Effect** | ***F* (df)** | **Significance** | **Partial Eta Squared** | ***t* (df)** | **Significance** |
| *All cell types* | | | | | |
| Line | 3.527 (1,12) | *p* = 0 .085 | 0.227 | 1.878 (12) | *p* = 0.085 |
| Sub-region | 100.349 (2,11) | ***p* < .0005** | 0.948 | | |
| Line X Sub-region | 0.863 (2,11) | *p* = 0.448 | 0.136 | | |
| *Ramified* | | | | | |
| Line | 0.829 (1,12) | *p* = 0.380 | 0.065 | 0.911 (12) | *p* = 0.380 |
| Sub-region | 116.704 (2,11) | ***p* < .0005** | 0.955 | | |
| Line X Sub-region | 0.922 (2,11) | *p* = 0.427 | 0.144 | | |
| *Reactive* | | | | | |
| Line | 1.158 (1,12) | *p* = 0.303 | 0.088 | 1.076 (12) | *p* = 0.303 |
| Sub-region | 14.173 (2,11) | ***p* = 0.001** | 0.720 | | |
| Line X Sub-region | 0.060 (2,11) | *p* = 0.942 | 0.011 | | |
| *Amoeboid* | | | | | |
| Line | 0.968 (1,12) | *p* = 0.345 | 0.075 | 0.984 (12) | *p* = 0.345 |
| Sub-region | 5.547 (2,11) | ***p* = 0.022** | 0.502 | | |
| Line X Sub-region | 2.257 (2,11) | *p* = 0.151 | 0.291 | | |

**Table S7. Total number of cells (per subject) included in area estimates using Nucleator probe**

| | CA1 | | CA3 | | DG | |
|---|---|---|---|---|---|---|
| | **bHRs** | **bLRs** | **bHRs** | **bLRs** | **bHRs** | **bLRs** |
| *Ramified* | 189.570 ± 6.969 | 212.860 ± 8.031 | 118.140 ± 6.749 | 121.000 ± 7.537 | 187.000 ± 9.097 | 180.570 ± 11.376 |
| *Reactive* | 14.290 ± 2.427 | 17.710 ± 3.428 | 7.860 ± 1.438 | 11.570 ± 1.875 | 20.570 ± 4.423 | 25.570 ± 4.418 |
| *Amoeboid* | 0.860 ± 0.340 | 2.570 ± 0.649 | 0.710 ± 0.286 | 1.000 ± 0.535 | 2.290 ± 0.747 | 1.570 ± 0.429 |

### Table S8. Statistics from soma and process area estimate analyses (Nucleator probe)

| | Mixed ANOVAs (line x sub-region) | | | | Independent t-tests (bHR v. bLR) | |
|---|---|---|---|---|---|---|
| | **Effect** | ***F* (df)** | **Significance** | **Partial Eta Squared** | ***t* (df)** | **Significance** |
| **Soma area** | | | | | | |
| *Ramified cells* | | | | | | |
| | Line | 2.465 (1,12) | *p* = 0 .142 | 0.170 | 1.524 (12) | *p* = 0 .153 |
| | Sub-region | 37.662 (2,11) | ***p* < 0.001** | 0.873 | | |
| | Line X Sub-region | 0.192 (2,11) | *p* = 0.828 | 0.034 | | |
| *Reactive cells* | | | | | | |
| | Line | 0.005 (1,12) | *p* = 0 .947 | 0.000 | 0.082 (12) | *p* = 0 .936 |
| | Sub-region | 3.387 (2,11) | *p* = 0.071 | 0.381 | | |
| | Line X Sub-region | 3.109 (2,11) | *p* = 0.085 | 0.361 | | |
| *Amoeboid cells* | | | | | | |
| | Line | 1.160 (1,12) | *p* = 0 .394 | 0.367 | 1.247 (12) | *p* = 0 .236 |
| | Sub-region | 1.634 (2,11) | *p* = 0.484 | 0.766 | | |
| | Line X Sub-region | 1.337 (2,11) | *p* = 0.522 | 0.728 | | |
| **Territory area** | | | | | | |
| *Ramified cells* | | | | | | |
| | Line | 59.772 (1,12) | ***p* < 0.001** | 0.833 | 7.709 (12) | ***p* < 0.001** |
| | Sub-region | 1.114 (2,11) | *p* = 0.363 | 0.168 | | |
| | Line X Sub-region | 1.939 (2,11) | *p* = 0.190 | 0.261 | | |
| *Reactive cells* | | | | | | |
| | Line | 13.36 (1,12) | ***p* = 0.003** | 0.527 | 3.843 (12) | ***p* = 0.002** |
| | Sub-region | 0.003 (2,11) | *p* = 0.997 | 0.001 | | |
| | Line X Sub-region | 0.112 (2,11) | *p* = 0.895 | 0.002 | | |
| *Amoeboid cells* | | | | | | |
| | Line | 0.863 (1,12) | *p* = 0.606 | 0.633 | 0.472 (12) | *p* = 0.645 |
| | Sub-region | 0.556 (2,11) | *p* = 0.688 | 0.526 | | |
| | Line X Sub-region | 0.067 (2,11) | *p* = 0.820 | 0.032 | | |

### Table S9. Statistics from detailed analyses of microglia morphology

| Anatomical measure | Interaction effects (Line x Sub-region) | | | | Independent t-tests (bHR v. bLR) | |
|---|---|---|---|---|---|---|
| | F (df) | Significance | Partial Eta Squared | | t (df) | Significance |
| Number of processes | 2.550 (2,73) | $p$ = 0.085 | 0.065 | *CA1* | 1.446 (74) | $p$ = 0.152 |
| | | | | *CA3* | 1.185 (74) | $p$ = 0.240 |
| | | | | *DG* | 1.143 (74) | $p$ = 0.257 |
| Process length | 7.637 (2,73) | **$p$ = 0.001** | 0.173 | *CA1* | 3.322 (74) | **$p$ = 0.001** |
| | | | | *CA3* | 5.765 (74) | **$p$ < 0.001** |
| | | | | *DG* | 1.131 (74) | $p$ = 0.262 |
| Process surface area | 5.964 (2,73) | **$p$ = 0.004** | 0.140 | *CA1* | 4.116 (74) | **$p$ < 0.001** |
| | | | | *CA3* | 6.686 (74) | **$p$ < 0.001** |
| | | | | *DG* | 2.474 (74) | **$p$ = 0.016** |
| Number of branches | 5.258 (2,73) | **$p$ = 0.007** | 0.126 | *CA1* | 6.267 (74) | **$p$ < 0.001** |
| | | | | *CA3* | 7.012 (74) | **$p$ < 0.001** |
| | | | | *DG* | 4.022 (74) | **$p$ < 0.001** |
| Convex hull area | 1.991 (2,73) | $p$ = 0.144 | 0.052 | *CA1* | 3.909 (74) | **$p$ < 0.001** |
| | | | | *CA3* | 4.869 (74) | **$p$ < 0.001** |
| | | | | *DG* | 1.733 (74) | $p$ = 0.087 |
| Soma area | 1.022 (2,73) | $p$ = 0.365 | 0.027 | *CA1* | 2.711 (74) | **$p$ = 0.008** |
| | | | | *CA3* | 3.043 (74) | **$p$ = 0.003** |
| | | | | *DG* | 1.311 (74) | $p$ = 0.194 |
